# COVID-19 Knowledge Graph: a computable, multi-modal, cause-and-effect knowledge model of COVID-19 pathophysiology

**DOI:** 10.1101/2020.04.14.040667

**Authors:** Daniel Domingo-Fernández, Shounak Baksi, Bruce Schultz, Yojana Gadiya, Reagon Karki, Tamara Raschka, Christian Ebeling, Martin Hofmann-Apitius, Alpha Tom Kodamullil

## Abstract

**Summary:** The past few weeks have witnessed a worldwide mobilization of the research community in response to the novel coronavirus (COVID-19). This global response has led to a burst of publications on the pathophysiology of the virus, yet without coordinated efforts to organize this knowledge, it can remain hidden away from individual research groups. By extracting and formalizing this knowledge in a structured and computable form, as in the form of a knowledge graph, researchers can readily reason and analyze this information on a much larger scale. Here, we present the COVID-19 Knowledge Graph, an expansive cause-and-effect network constructed from scientific literature on the new coronavirus that aims to provide a comprehensive view of its pathophysiology. To make this resource available to the research community and facilitate its exploration and analysis, we also implemented a web application and released the KG in multiple standard formats.

**Availability:** The COVID-19 Knowledge Graph is publicly available under CC-0 license at https://github.com/covid19kg and https://bikmi.covid19-knowledgespace.de.

**Supplementary information:** Supplementary data are available online.

## 1. Introduction

The COVID-19 crisis has prompted a response of the scientific community that is unparalleled in history. Research organizations have dedicated their entire workforce to combat the pandemic. Tens of thousands of researchers in hundreds of universities, governmental laboratories, and industrial research departments have entirely focused their efforts on understanding the virus pathophysiology, finding drugs that interfere with its life cycle and developing immunization strategies for future vaccines (Chahrour *et al.*, 2020).

While the steep increase in research activities in the COVID-19 context has led to an unprecedented increase of scientific publications, it becomes challenging to identify genuine novel findings and discern them from those that are already known. The process of discriminating “known knowns” from “unknown knowns” can be supported by knowledge graphs (KGs), as they provide a means to capture, represent and formalize structured information (Nelson *et al.*, 2019). Furthermore, although these KGs were originally developed to describe interactions between entities, they are complemented by a broad range of algorithms that have been proven to partially automate the process of knowledge discovery (Cowen *et al.*, 2017; Humayun *et al.*, 2020). Importantly, novel machine learning techniques can generate latent, low-dimensional representations of the KG which can then be utilized for downstream tasks such as clustering or classification (Hamilton *et al*., 2015).

In this paper, we present an approach to lay the foundation for a comprehensive KG in the context of COVID-19. Our work is complemented by a web application that enables users to comprehensively explore the information contained in the KG. To facilitate the ease of usage and interoperability of our KG, we have released its content in various standard formats in order to promote its adoption and enhancement by the scientific community.

## 2. Material and Methods

In this section, we outline the methodology used to: i) select the corpus, ii) generate the COVID-19 KG, and iii) develop the web application for exploring the KG.

### 2.1 Selection of Scientific Literature

For the creation of the KG, scientific literature related to COVID-19 was retrieved from open access and freely available journals **(Supplementary Information)**. This corpus was then filtered based on available information about potential drug targets for COVID-19, biological pathways in which the virus interferes in order to replicate in its human host, and information on the various viral proteins along with their functions. Finally, the articles were prioritized based on the level of information that could be captured in the modeling language used to build the KG.

### 2.2 Constructing the COVID-19 Knowledge Graph

Evidence text from the prioritized corpus was manually encoded in Biological Expression Language (BEL) as a triple (i.e., source node - relation - target node) including metadata about the nodes and their relationships as well as corresponding provenance and contextual information. BEL scripts generated from this curation work are freely available at https://github.com/covid19kg along with their network representations in several other standard formats (e.g., SIF, GraphML, and NDEx). By making this data available in multiple formats, we are seeking to facilitate the analysis of the KG with a broad range of methods/software as well as promote its integration into other biological databases and web services such as the one presented in the following subsection.

### 2.3 Web Application

In order to better aid the exploration and usage of the generated COVID-19 Knowledge Graph, a web application was developed using Biological Knowledge Miner (BiKMi), an in-house software package designed for exploring pathways and molecular interactions within a BEL-derived network. The front-end of the application was constructed using the Python Django web framework, while the back-end of the software is implemented using OrientDB, a multi-model database management system that allows for both relational and graph queries to be made against a database via its API **(Supplementary Information)**, which opens the avenue towards systematic comparison of different COVID models.

## 3. Results

We introduce a KG that comprises mechanistic information on COVID-19 published in 145 original research articles. In its current state, the COVID-19 KG incorporates 3954 nodes, covering 10 entity types (e.g., proteins, genes, chemicals, and biological processes) and 9484 relationships (e.g., increases, decreases, and association), forming a seamless interaction network **(Supplementary Information)**. Given the selected corpora, these cause-and-effect relations primarily denote host-pathogen interactions as well as comorbidities and symptoms associated with COVID-19. Furthermore, the KG contains molecular interactions related to host invasion (e.g., spike glycoprotein and its interaction with the host via receptor ACE2) and the effects of the downstream inflammatory, cell survival, and apoptosis signaling pathways.

A key aspect of the COVID-19 KG is in its large coverage of drug-target interactions along with the biological processes, genes and proteins associated with the novel coronavirus. We have identified over 300 candidate drugs currently being investigated in the context of COVID-19 **(Supplementary Information)**, including proposed repurposing candidates and drugs under clinical trial.

Along with the KG, we implemented a web application (https://bikmi.covid19-knowledgespace.de) for querying, browsing, and navigating the KG **(Figure 1)**. The visualization enables users to explore and query the network (e.g., filtering nodes or edges, or calculating paths between nodes of interest). Additionally, it enables users to upload *omics* data and validate its signals against the knowledge contained in the network. To demonstrate this feature, the web application is loaded with the transcriptomics experiments conducted by Blanco-Melo *et al.* (2020).

**Figure 1.**
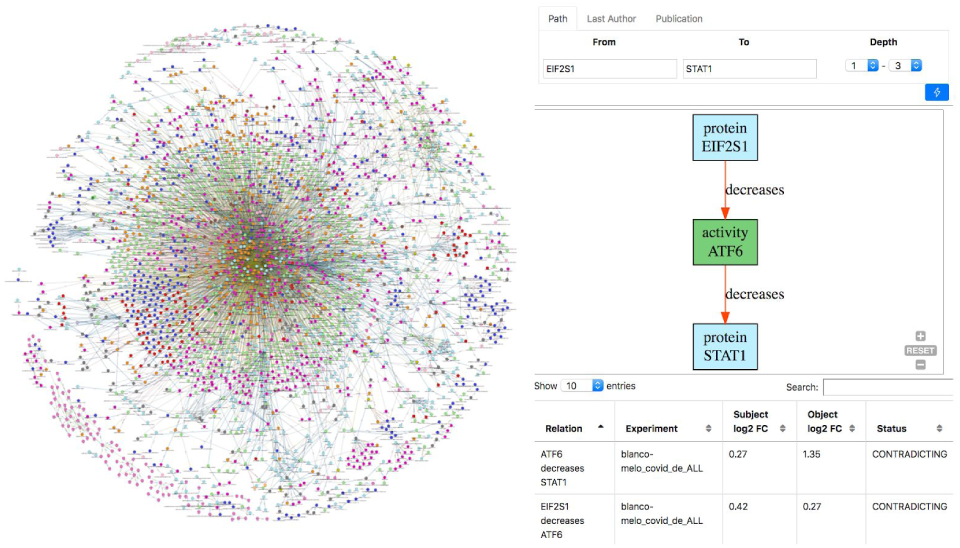
(Left) Visualization of the COVID-19 KG in BiKMi. (Right) Querying paths between two nodes and verifying their consistency with transcriptomics data.

## 4. Discussion

The novel coronavirus has motivated a profound response by the scientific community and has led to the rapid publishing of COVID-19 research. As an attempt to organize and formally represent the most current knowledge of the virus, we have introduced a KG comprising mechanistic information around COVID-19 biology and pathophysiology. The presented KG provides a comprehensive overview of relevant viral protein interactions and their downstream molecular mechanisms. Additionally, it also includes the vast majority of potential drug-targets as well as clinical manifestations associated with comorbidities and symptoms. Given the biological complexity and the sparse information we currently have on the pathophysiology of the virus, mechanistic knowledge contained in the KG could be promising for the discovery of as of yet hidden interactions. The COVID-19 KG presented here is part of a bigger ecosystem that integrates disease maps with three of the largest pathway databases (Domingo Fernandez *et al*., 2019).

Not only do we provide a web application to make the content accessible to the research community, but we also have released the KG in a variety of standard formats. In doing so, we aim to foster an exchange of information across similar integration approaches (Waagmeester *et al.*, 2020) as well as to facilitate its analytic use on both knowledge- and data-driven methods. Finally, we plan to make future releases of the KG to ensure the most up-to-date content as well as to benefit from its integration and crosstalk with other similar activities (i.e., #covidpathways).

