## Supplementary Information for "COVID-19 Knowledge Graph: a computable, multi-modal, cause-and-effect knowledge model of COVID-19 pathophysiology"

### Additional File

#### Outline

1. Selection and Prioritization of the Knowledge Graph Corpus
2. Knowledge Graph Content
3. Applications
4. Web Application

#### Supplementary Text

##### 1. Selection and Prioritization of the Knowledge Graph Corpus

As the pandemic grew, several open access and freely available resources were made available to query about COVID-19. We curated several of these open source resources as well as scientific corpora for the creation of the knowledge graph (KG) corpus. Prior to curating all the available research articles, a pre-selection guideline was established amongst curators. These guidelines involved both informational as well as programmatic aspects. For informational aspects, we ensured that the articles selected would contain information underlying molecular mechanisms, etiology and potential drug targets for COVID-19. Due to its novelty, apart from the specific focus on SARS-CoV-2, information about other coronaviruses such as SARS-CoV was also taken into account. The preferred articles based on the criteria outlined in the manuscript were encoded in BEL, the format used to generate the KG. BEL involves encoding mechanistic information such as protein-protein interactions, observed correlations between phenotypes and molecules, or effect of drugs on a given target. Therefore, the articles were also selected taking into consideration this aspect (i.e., BEL encodable articles).

Taking into consideration one of the largest and freely available data source dump, PubMed and EuropePMC were our primary choices for the generation of corpus. Hence, we queried them with general MeSH terms such as "coronavirus" and "COVID-19" followed by specific queries such as ("pi3k and viral replication") and ("pi3k and sars"). As the wave of COVID-19 research advanced, a number of other corpuses specific for COVID-19 were becoming popular. Thus, we used these specific corpuses to further enrich our KG. With this aim, LitCovid was one of the external freely available corpuses used. It is a COVID-19 specific curated literature hub from NCBI which provides publications under several headings such as mechanism, transmission, diagnosis, treatment and prevention. We selected review articles from the treatment section that provided insights about the biology of the virus or potential treatment or both. Since only a small number of these articles could be converted to BEL, we shifted to another corpus named Targeting2019-nCoV, created by GHDDI and made availalble at https://ghddi-ailab.github.io/Targeting2019-nCoV. Upon the analysis of KG at this point, we understood that there were some missing links with respect to host-pathogen interactions and molecular mechanisms. Hence, a third corpus provided by University of Luxembourg (https://covid.pages.uni.lu) at https://www.zotero.org/groups/2471739/covid-19- graph-curation was considered. Thus in summary, along with articles from PubMed and EuropePMC, a number of additional COVID-19 specific corpuses like LitCovid (by NCBI), Targeting2019-nCoV (by GHDDI) and University of Luxembourg were source for creation of KGs.

To demonstrate how other resources could be integrated into the KG (e.g., #covidpathways), we made use of these published resources. With the focus on drug-target information, we retrieved drug-target information which were either drug-viral protein interactions or drug-human protein interactions from DrugBank (version 5.1.5) (Wishart *et al*., 2008). Additionally, the COVID-19 interactome published by Gordon *et al.* (2020) on PathwayCommons (Cerami *et al.*, 2011) was merged with the current knowledge graph. However, these two resources have not been included in the statistics since they have not been manually curated but imported.

In the end, a total of 145 full text reviews, along with data from two databases (i.e., PathwayCommons and DrugBank), were selected for curation and creation of the COVID-19 Knowledge Graph (https://github.com/covid19kg/covid19kg/blob/master/supplement/summary.csv). This and future releases of the KG will be archived using Zenodo (https://doi.org/10.5281/zenodo.3748950).

#### 2. Knowledge Graph Content

In this section, we outline the content of the KG which is also summarized in the Jupyter Notebook available in the following GitHub repository link https://github.com/covid19kg/Analysis/blob/master/notebooks/tutorial.ipynb. **Supplement Figure 1** shows the different types of nodes (left) and edges (right) on the first release on the KG. The most common nodes are complexes, proteins, biological complexes, and abundance due to the strong focus of the KG towards covering the molecular aspects of the virus. Statistically speaking, we identified 684 biological processes and 1,132 genes and proteins that were associated with the virus. Regarding relations, the most common ones are positive correlations and associations (largely covering relations between phenotypes and other clinical observations) and mechanistic relations such as increases and decreases. **Supplement Figure 2** shows the different database information present in the KG. As we can infer, most of the information contained within the KG is about proteins and biological processes followed by chemicals or drugs.

Although the complete process of virus-host interaction and its physiological consequences at the molecular level is yet to be explored, our knowledge model has a strong focus on available repurposed as well as custom tested drug information and the effect on their targets. Out of the drugs found in our KG, the most promising drugs based on a number of positive outcomes from different studies are lopinavir, ritonavir, remdesivir, chloroquine, and oseltamivir (Guo *et al.*,2020, Cunningham *et al.* 2020, Rismanbaf 2020, Weston *et al.* 2020). Antiviral lopinavir in combination with ritonavir is an approved drug for HIV infection and currently under investigation for COVID-19 (Li *et al*. 2012). Remdesivir is an adenosine triphosphate analog under investigation for Ebola and different coronaviruses (Warren *et al.* 2016). Chloroquine or hydroxychloroquine is a well known anti-malarial drug that inhibits the conversion of heme to hemazoin in malarial trophozoites (Slater *et al.* 1992, Coronado *et al.* 2014). Oseltamivir is another antiviral drug and antiviral neuraminidase inhibitor that showed positive outcomes in some COVID-19 cases (Cooper *et al.* 2003, Cunningham *et al.* 2020). In summary, our knowledge graph provides an overview of the viral-interaction and downstream molecular mechanisms inside the host, potential drug-targets as well as clinical manifestations in terms of associated morbidities and symptoms.

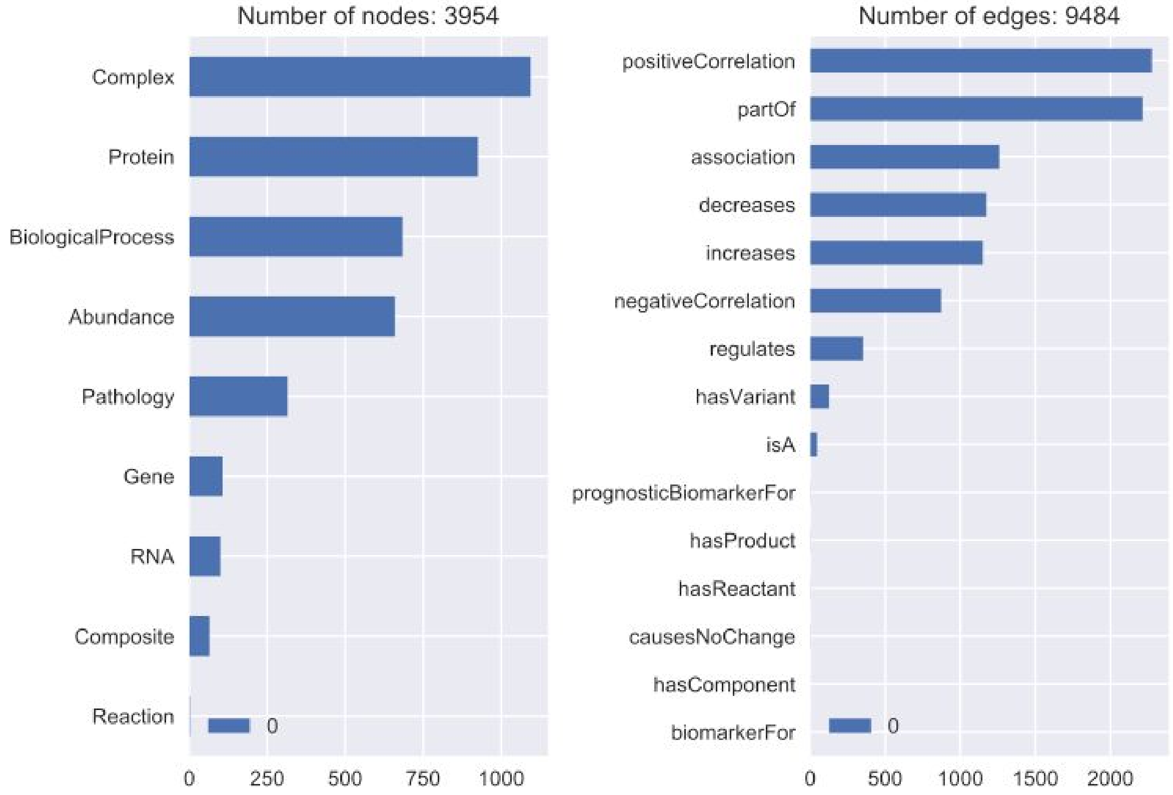


**Supplement Figure 1.** Distribution of the different nodes (left) and edges (right) encoded in the COVID-19 Knowledge Graph.

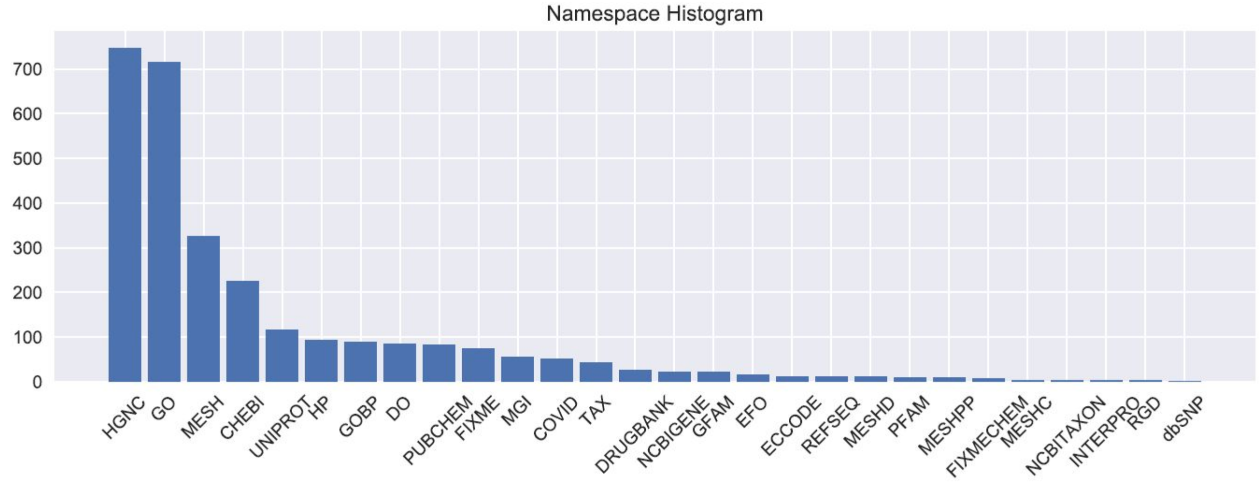


**Supplementary Figure 2.** Distribution of namespaces used to encode nodes in the COVID-19 Knowledge Graph.

To illustrate the current size of the KG, we generated **Supplementary Figure 3**. However, we do not recommend exploring the full network directly on web applications that rely on JavaScript. Instead, we commend the use of GraphML files and recommend the use of dedicated software such as Cytoscape (https://cytoscape.org/). As expected, the KG containing all the above-mentioned information is densely connected. On the contrary, there still seems to be a lot of unknown information required to enrich the KG more, so as to find the missing links between the isolated nodes and fully connected ones.

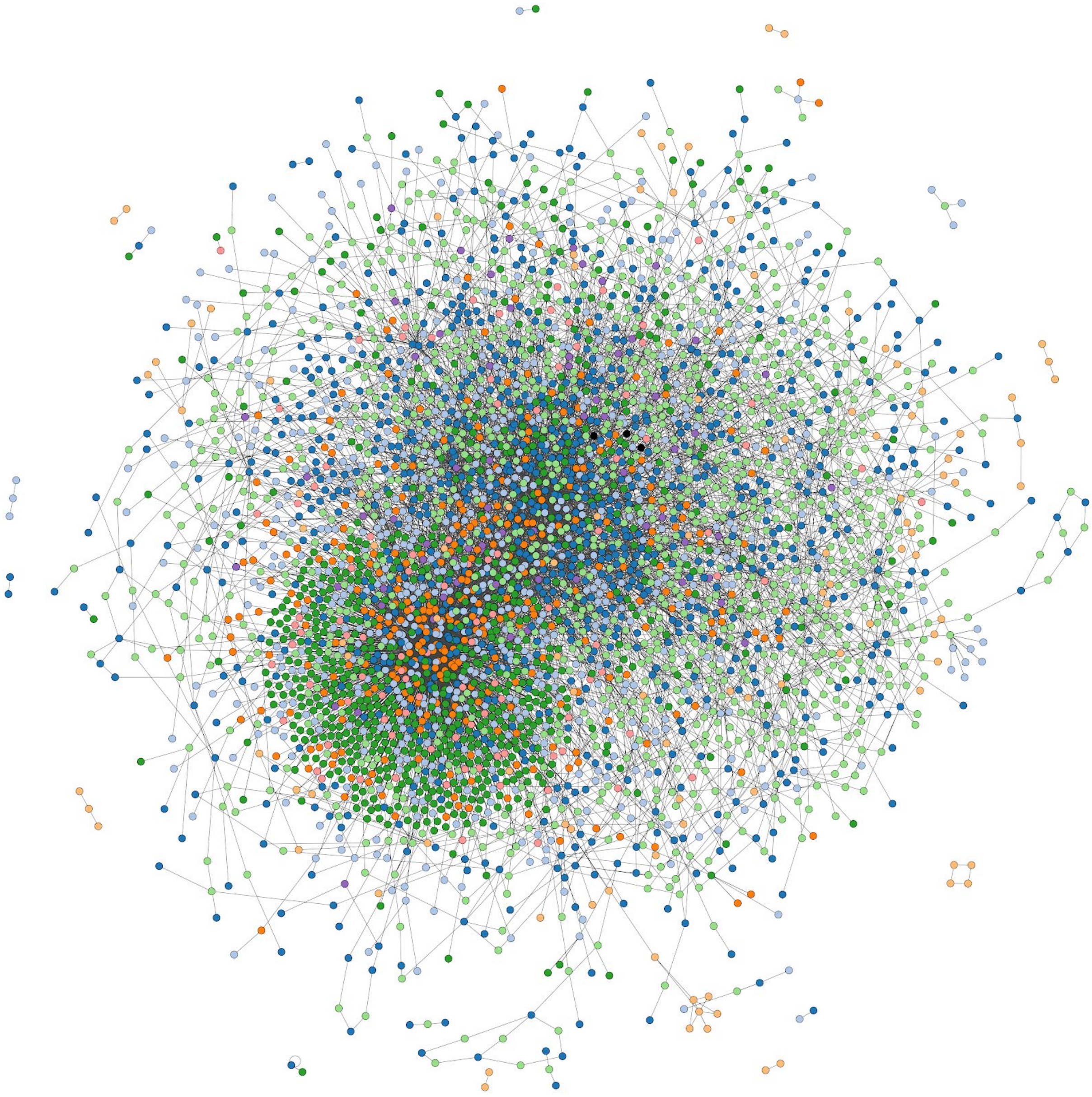

**Supplementary Figure 3.** A quick glance at the amount of information stored in the COVID-19 Knowledge Graph.

### 3. Applications

#### 3.1. Gene Expression Analysis

A differential gene expression analysis was done on raw read counts provided through GEO series GSE147507 (Blanco-Melo *et al.,* 2020). The analysis was performed with the help of DESeq2 [v1.22.2] (Love *et al.* 2014) in R. After quality control and removing transcripts with no counts, the differential gene expression analysis was done using different conditions as contrast, resulting in five interpretable comparisons, namely 'mock' vs 'SARS-CoV-2 infected' 1) for all cell types, 2) for NHBE cells, 3) for A549 cells, 4) for A549 cells with vector expressing human ACE2, and 5) for

Calu-3 cells. Differentially expressed genes with an adjusted p-value $< 0.05$ and an absolute log fold change $> 1$, were chosen for further analysis.

##### 3.2. Pathway Enrichment Analysis on the COVID-19 KG

As an illustration of a simple analysis, we conducted pathway enrichment using the human genes presented in the COVID-19 KG using a one-sided Fisher's exact test (Fisher, 1992) against three pathway databases (i.e., KEGG, Reactome, and WikiPathways) (Kanehisa *et al.,* 2016; Fabregat *et al.*, 2018; Slenter *et al.*, 2017). We consider a pathway to be significantly enriched if its $q$-value is smaller than 0.05 after applying multiple hypothesis testing correction with the Benjamini–Yekutieli method under dependency (Benjamini and Yekutieli, 2001). This analysis is presented at the following notebook: https://github.com/covid19kg/Analysis/blob/master/notebooks/enrichment_analysis.ipynb.

#### 4. Web Application

In this section, we provide a brief overview of our web application designed specifically for the ease of interoperability and data sharing of COVID-19 Knowledge Graph amongst interested researchers. This web application uses two key components: the open-source Django web framework (https://www.djangoproject.com/) to render the front-end of the service, and OrientDB (https://orientdb.com/) for the back-end to both house the network itself and run the API functions for the requested information. The following subsections outline the front-end and back-end development related to the web application.

##### 4.1. Front End

Django was written using Python and enables rapid web application development due to its ease-of-use and built-in libraries while also ensuring compatibility with most of the major operating systems. The front-end was designed as a single-page application (SPA) in which instead of loading a new web page for each available feature of the application (multi-page application), a single web page is dynamically rewritten with the desired content. This was chosen to better provide users with a faster and more responsive experience.

##### 4.2. Back-End

In order to present the relevant information to the user, Django must first collect the necessary data from the back-end of the web application. The OrientDB NoSQL database management system (DBMS) stores all of the curated BEL triples in a single database and generates a graph from the defined interactions. Because it is both a relational and graph database, OrientDB requires a query language with both table and graph query functionality, therefore it created a modified SQL dialect that includes extensions which enable graph functionality. In addition to being able to query both the graph and tables directly, this DBMS also includes a RESTful API for which specific functions can be written to extract specific information from the database. These functions can be written using the modified SQL query language that OrientDB developed, Apache Groovy Language (https://groovy-lang.org/) or JavaScript (https://www.javascript.com/), but in all instances the RESTful API returns the requested data in the JSON format. The RESTful API functions used for our web application were constructed automatically using in-house software e(BE:L).

Furthermore, the OrientDB DBMS also has the added benefit of being able to order node and edge types in a hierarchical format such that a particular node or edge class can inherit properties from a parent class. This enables one to develop a defined schema that groups together similar classes using a superclass which can then be used for creating precise and well-defined queries that do not depend on listing individual node and edge types. For example, nodes representing proteins, RNA, and genes can inherit from a superclass called "genetic_molecules" and this superclass can be used to query against all 3 node types at once.
